# Complete genome sequence and analysis of nine Egyptian females with clinical information from different geographic regions in Egypt

**DOI:** 10.1101/2020.03.10.985317

**Authors:** Mahmoud ElHefnawi, Elsayed Hegazy, Asmaa ElFiky, Yeonsu Jeon, Sungwon Jeon, Jong Bhak, Fateheya Mohamed Metwally, Sumio Sugano, Terumi Horiuchi, Abe Kazumi, Asta Blazyte

## Abstract

Egyptians are at a crossroad between Africa and Eurasia, providing useful genomic resources for analyzing both genetic and environmental factors for future personalized medicine. Two personal Egyptian whole genomes have been published previously and here nine female whole genome sequences with clinical information have been added to expand the genomic resource of Egyptian personal genomes. Here we report the analysis of whole genomes of nine Egyptian females from different regions using Illumina short-read sequencers. At 30x sequencing coverage, we identified 12 SNPs that were shared in most of the subjects associated with obesity which are concordant with their clinical diagnosis. Also, we found mtDNA mutation A4282G is common in all the samples and this is associated with chronic progressive external ophthalmoplegia (CPEO). Haplogroup and Admixture analyses revealed that most Egyptian samples are close to the other north Mediterranean, Middle Eastern, and European, respectively, possibly reflecting the into-Africa influx of human migration. In conclusion, we present whole-genome sequences of nine Egyptian females with personal clinical information that cover the diverse regions of Egypt. Although limited in sample size, the whole genomes data provides possible geno-phenotype candidate markers that are relevant to the region’s diseases.

## 1. Introduction

Next-generation sequencing (NGS) technology is a powerful approach enabling an efficient study of a large scale genetic variants [1,2,3,4]. Whole genome sequencing (WGS) by NGS can cover intronic areas that may contain rare and common deleterious mutations, that cannot be captured by whole exome sequencing (WES) that usually facilitates a deeper coverage of coding regions important for protein function analyses [5]. NGS is also adopted to investigate complex diseases such as obesity and its manifestations such as insulin resistance, impaired glucose tolerance, and dyslipidemia. For instance, Nordang *et al.* [6] applied NGS technology to target exons and exon-intron junctions in the five obesity-linked genes (*LEP, LEPR, MC4R, PCSK1* and *POMC*) for a Norwegian-based cohort composed of patients with morbid obesity and normal weight controls. They found four variants in *MC4R* gene that were classified as pathogenic or likely pathogenic. Before that, Genome-Wide Association Studies (GWAS) have identified several variants associated with obesity and BMI in the *FTO* gene [7,8]. Environmental factors such as socioeconomic status and lifestyle, can complement the effect of the obesogenic factors [9,10,11,12]. However, the response to the environmental factors can also be ethnic-group specific which makes GWAS difficult to interpret. For example, Caucasians with different genetic backgrounds, but living in a similar obesogenic environment are less susceptible to developing obesity (∼32 %) or T2D (∼8 %) when compared to Pima Indians who live in Arizona where obesity and T2D in those populations were ∼64% and ∼30%, respectively [13,14].

As for Egyptians, obesity studies have exceptional relevance as it is one of the nations with the steepest obesity rate in a 33 yearlong study (1980-2013) [15]. A study published in 2016 reported more than 30 percent of the Egyptian population to be obese with nearly half of all females being obese (46%) which is nearly two times the magnitude of this issue in Egyptian males (22%) [16]. The majority (66%) of Egyptian women is overweight, in fact, Egyptian women of reproductive age in just 10 years showed an average increase in BMI by 2.21 kg / m ^2^ [17]. A recent PCR-based study came to a conclusion that *LEPR* Gln223Arg, *UCP2* G 866 A and *INSR* exon 17 polymorphisms are linked to obesity in Egyptians and some of the risk variants are indeed sex biased [18]. In our previous work the two Egyptian male whole genomes designated as (EGP1 and EGP2) from Delta and Said regions, respectively, shared the SNPs Asn985Tyr in *RP1* and Thr55Ala in *FABP2* which are associated with an increased level of cholesterol and triglycerides, possibly related to obesity [19].

## 2. Materials and methods

### 2.1. Study group and ethical approval

A total of nine Egyptian females were recruited for this study, all aged 41±14.35 years. The subjects (EGY) were enrolled by National Research Center’s outpatient clinic and approved by the Medical Research Ethics Committee of the National Research Centre –Egypt (MREC, NRC) with approval number (13177) and with 1964 Helsinki Declaration and its later amendments. Signed informed consents were obtained from the participants in this study. A clinical examination was conducted for each of them; weight (weight), height (Ht) and waist circumference (WC) were measured. Body mass index (BMI) was calculated according to WHO [20]. Blood pressure was measured for each individual. Five ml of whole blood was centrifuged at 12,000 rpm for 10 minutes using (Heraeus Labofuge 400R) centrifuge to separate serum and stored at −80C° until analysis.

### 2.2. Biochemical Analyses

Total cholesterol, the levels of triglyceride and High-Density lipoprotein cholesterol (HDL-c) were assessed using an enzymatic colorimetric method by the commercial kits supplied by Centronic GmbH (Wartenberg, Germany). Serum low density lipoprotein cholesterol (LDL-C) was estimated using Friedwald’s formula.

Fasting serum glucose was assessed using commercial kit (Spinreact,Spain), Fasting serum insulin was estimated by human insulin enzyme immunoassay test kit (Immunospec, USA). Insulin resistance was expressed as homeostasis model assessment of insulin resistance (HOMA-IR) higher than 2.5 [21].

### 2.3. Sample collection, DNA isolation and whole genome sequencing

Genomic DNA was extracted from blood with the GeneJET (Whole Blood) genomic DNA purification Kit (Thermo Scientific, USA), according to manufacturer’s protocol. DNA Library preparation was carried out according to the manufacturer’s instructions for sequencing on HiSeq2000 (Illumina, San Diego, CA, USA). For demultiplexing and conversion to FASTQ format, CASAVA 1.8.2 (Illumina) was used.

### 2.4. Bioinformatics analysis

#### 2.4.1. Alignment of reads to reference

The sequences were mapped to the human reference genome (GRCh38 /hg 38) BWA-MEM v.0.7.15 with default options [22]. The SAM files were converted to BAM files using samtools 0.1.19 [23]. GATK v.3.5.0 [24] was used for variant calling. Specifically, GATK v.3.5.0 UnifiedGenotyper with options -mbq 30 -stand_emit_conf 50 -stand_call_conf 50.

#### 2.4.2. Annotation and functional analysis of variants

The variants were annotated using Annovar v.2018Apr16 [25] and SnpEff tool v4.3i [26]. CRAVAT tool v.4 [27] was used for predicting mutation impact. It employs Variant Effect Scoring Tool (VEST) which classified mutations as pathogenic or benign. Furthermore, ClinVar database was used [28] to predict the pathogenic mutations.

#### 2.4.3. Identification of mitochondrial DNA haplogroups

The paired-end reads aligned to hg19 mitochondrial sequence were realigned to rCRS (Revised Cambridge Reference Sequence [29] and then the variants were used to call haplogroups using HaploGrep software [30].

#### 2.4.4 Admixture analysis

To visualize the heterogeneity of the Egyptian lineages we employed ADMIXTURE (v1.3.0) program [31]. We used a total of 80 human origin SNP panel (HOSP) [32] contemporary representative genomes from Europe, West Asia, and various parts of Africa (four per population). The panel was merged with the Egyptian samples and pruned using PLINK (v.1.90) [33] with the option ‘--indep-pairwise 200 25 0.4’. We explored the composition of the artificial ancestral populations (*K*) from two to five to provide the glimpse into the increasing Egyptian genomic complexity as *K* increases (supplementary figure S1), and chose *K* = 4 for the final interpretation because it showed a gradual transition of ancestral component proportions from European and West Asian populations to North African as predicted by geographic location (supplementary figure S2). Out of the 9 Egyptian samples, sample EGY3 was excluded due to blood relations with another sample (EGY4) in the dataset to avoid bias.

## 3. Results

### 3.1. Clinical characteristics of the study group

The clinical characteristics of each subject are represented in supplementary Table S1. The mean value of BMI was 32.1±11.28; ranged from (18-45.9), where five cases (aged 44-58 years old) were obese with BMI greater than 30 kg / m ^2^. Four subjects (aged 20-35 years old) were classified as lean with BMI less than 25 kg / m ^2^. The mean level of total cholesterol among the research subjects was 192.7±51.5. Three cases of hyperlipidemia (aged 29-58 years old) were observed; in one of the cases coinciding with hypertension. Regarding HOMA-IR, the mean value was 2.3±1.59 and ranged from 0.9 to 5.08. Two out of the nine subjects (age 50 and 56 years old) were diagnosed with diabetes, and one had HOMA-IR 4.5 but without any confirmed diagnosis. Three out of the nine subjects (EGY3 age 35, EGY4 age 20, and EGY5 age 25 years old) weren’t clinically diagnosed with any metabolic syndrome.

### 3.2. Genome sequencing and candidate gene identification

The nine Egyptian females represent different geographic regions in Egypt (supplementary Figure S2). The workflow of the sequence pre-processing, variant calling, annotation and analysis is summarized in eight steps (supplementary Figure S3.). We used an Illumina HiSeq 2500 short-read sequencing and generated approximately 3,099,0381,190 reads of 100 bps length that were aligned to the human reference genome GRCh38/hg 38, resulting in the average depth of coverage of 30x. The basic sequence data are summarized in supplementary Table S2

From our data, we identified genes associated with obesity, T2D, and MS using the CRAVAT v.4 server and listed the ones with significant association (supplementary Table S3). Among them, we identified five candidate genes for obesity in the Egyptians; *CRHR1, VCP, NOTCH3, RHBG*, and *TMEM63A*. Variations in these genes were common within the subjects with BMI> 30 kg/m^2^. *CRHR1* and *TMEM63A* were associated with T2D-diagnosed subjects. Also, *NUP88, CRHR1*, and *WISP1* were associated with high cholesterol trait. The gene *CRHR1* variations were found in both high cholesterol and T2D patients and therefore this gene is the most feasible obesity and obesity-related disease risk factor we can suggest based on these nine Egyptian females studied.

### 3.3. Variants analysis

We used CRAVAT v.4 server to identify the most significant SNPs in genes using ClinVar and variant effect scoring tool (VEST) score, [34] as well as phenotypes reported in the GWAS catalogue [35] which resulted in a total of 12 SNPs related to metabolic issues and obesity shared among the subjects. Some of them are associated with the pathological conditions confirmed in the Egyptian (EGY) subjects (supplementary Table S4). Among them, we found rs3733402, Ser143Asn in *KLKB1* that are associated with obesity according to GWAS catalog, and rs351855, Gly388Arg in *FGFR4* that are associated with Waist-to-hip ratio adjusted for body mass index [36].

In the prehypertensive and hypertensive subjects (systolic and diastolic 130/90 &150/100) with high cholesterol, we found Asp131Glu in C1GALT1C1 (rs17261572, T>A), rs10065172, Leu105 in *IRGM* and rs1805010, Ile75Val in *IL4R* while in the prediabetic with HOMA-IR >2.5 and diabetic participants, we found the following variants: rs1804495, Leu303Phe in *SERPINA7*, rs1800450, Gly54Asp in *MBL2*, rs1566734, Gln276Pro in *PTPRJ*, and rs61752717, Met694Val in *MEFV*.

### 3.4. Mitochondrial DNA analysis

The mitochondrial DNA (mtDNA) haplogroups were identified using and MitoSuite 1.0.9 and validated using Haplogrep 2.0 tools (Table 1) and showed diverse maternal ancestral backgrounds associated with Africa, Middle East, Europe or broadly - Mediterranean. The sample EGY1 and EGY2 revealed haplogroups associated with Mediterranean regions, M1a1b found in the north Mediterranean [37,38,39,40] and T2f found in the European countries, respectively. The eastern Mediterranean region shares severe incidence of obesity with Egypt [41]showing a possible genetic linkage within these countries regarding metabolic profile. Whereas, EGY3 and EGY4 are related samples sharing haplogroup L2a1 and EGY5 had haplogroup M, in both cases, these haplogroups point to an African origin [42]. The strongest mtDNA ancestral connection to the Middle East was shown by EGY8 haplogroup H5 [43], however, such origin could be attributed to EGY6 (H27e) and EGY7 (U8b1a2b) as well EGY8 [44,45,46]. EGY7 haplogroup showed a very specific distribution within Jordan and Italy [47,48,49]; possibility of European or Middle Eastern admixture in this sample seems to be reflected in the ADMIXTURE analysis (supplementary Figure S1).man Haplogroup T1a7 found in the EGY9 is associated with very broad distribution within all of the geographic regions described regarding other EGY samples [50,51,52]. Moreover, mitochondrial genome of diabetic subjects and one of three cases of hyperlipidemia, EGY7, EGY9, and EGY2 contained T16189C mutation (Table 2) associated with insulin resistance and type 2 diabetes mellitus [53,54,55,56]. Among the variants that were not related to obesity, we found that A4282G (Table 2) associated with chronic progressive external ophthalmoplegia (CPEO) was present in all samples [57].

**Table 1:**
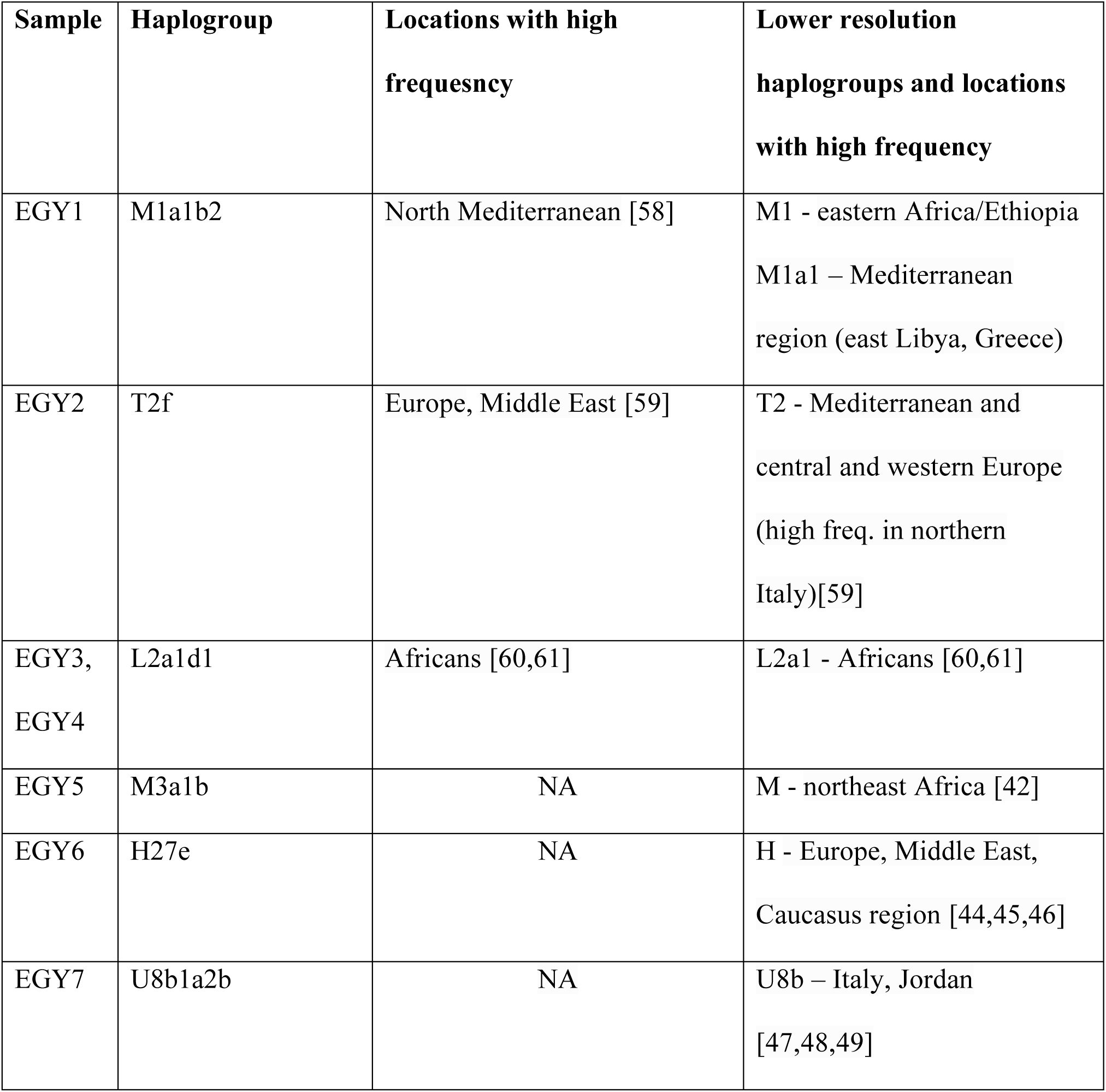
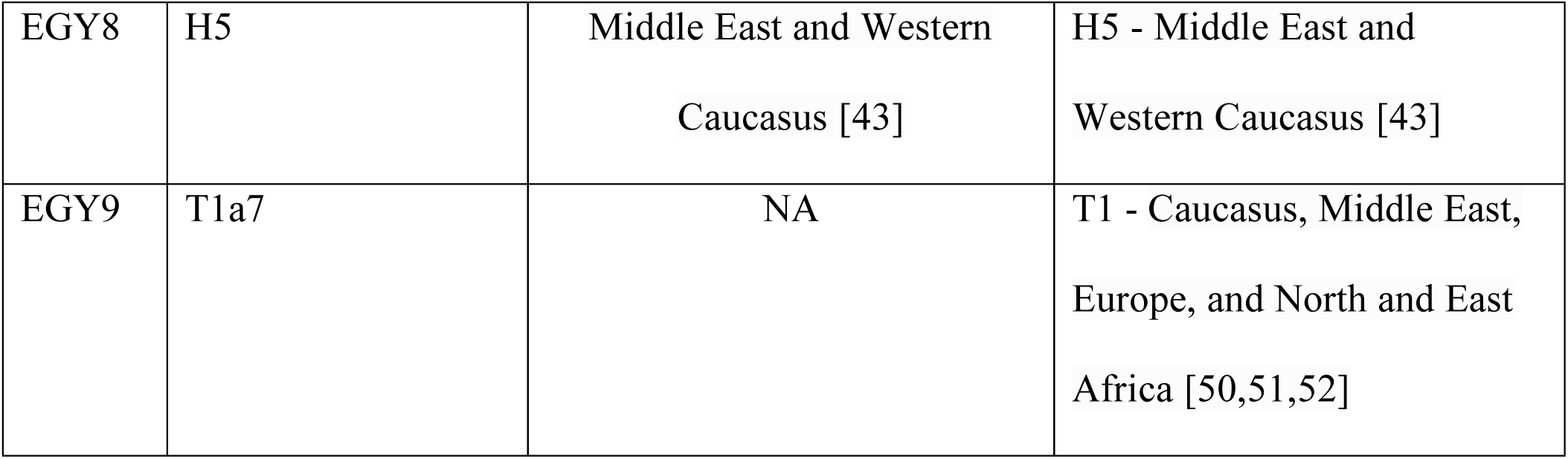
List of Haplogroups and locations where they are common regarding each sample.

**Table 2.** List of mitochondrial disease associated variants in each sample.

|  | Mutations (disease associated) |
| --- | --- |
| EGY1 | T195C, A4282G ,A8108G), A10398G ,G15043A, G16129A), A16183C and T16189C |
| EGY2 | T4216C ,A4282G), A4917G), A10398A) and G15928A |
| EGY3 | T195C, A4282G , A10398G T15784C , G16390A and T16519T |
| EGY4 | T195C , A4282G , A10398G (, T15784C, G16390A and 16519 |
| EGY5 | A4282G , A10398G and G15043A |
| EGY6 | A4282G , A10398A ), T16093C and T16519T |
| EGY7 | T195C ), G3316A , A4282G, G9055A , A10398A , A11467G, A12308G , G12372A,<br>G14831A , A16183C and T16189C |
| EGY8 | A4282G , A10398A ), C16192T and T16519T |
| EGY9 | T4216C , A4282G , A4917G , A10398A , G15928A and T16189C |

### 3.5. Admixture analysis

When *K* = 4 is chosen (Fig 1), the major ancestral genomic components in Egyptians are dark green and dark blue. While the dark green varied between 81.10% and 92.43% among the Egyptian samples, the dark blue composed from 7.56% to 18.48% of the artificial Egyptian ancestry. Similar genomic trend was shown when the Tunisians, Algerians, Saharawi and the Bedouin samples used, which points to a strong uniform North African genomic signature, despite of nomadic lifestyle practiced by some of these populations [62,63] which could have resulted in a high admixture rate. However, the components themselves seem to have originated from two main sources: the dark green from the Southern Europe as well as the Middle East and the dark blue mainly from the West Africa as it was found in the highest proportions in the Gambian, Essan, and Yoruba populations. Besides the European/Middle Eastern and West African components small amount and highly variable (from 0.00% to 1.05%) East-African-associated component (soft orange) (supplementary Table S5) was found in highest proportions in EGY9, EGY6 and EGY5 among the Egyptian samples. This component had the strongest association with the Hadza indigenous people, however, in smaller proportions was also found among North-Eastern (Ethiopia) and East-Central (Dinka) African populations potentially reflecting genetic drift across the African continent. Most of the Egyptian samples used, especially, EGY5 and EGY9 are closely resembled with Tunisian ancestral proportions while EGY7, EGY8 and EGY2 exhibited higher proportions of the green (Southern European/West Asian) component which were common among the Bedouin.

**Figure 1:**
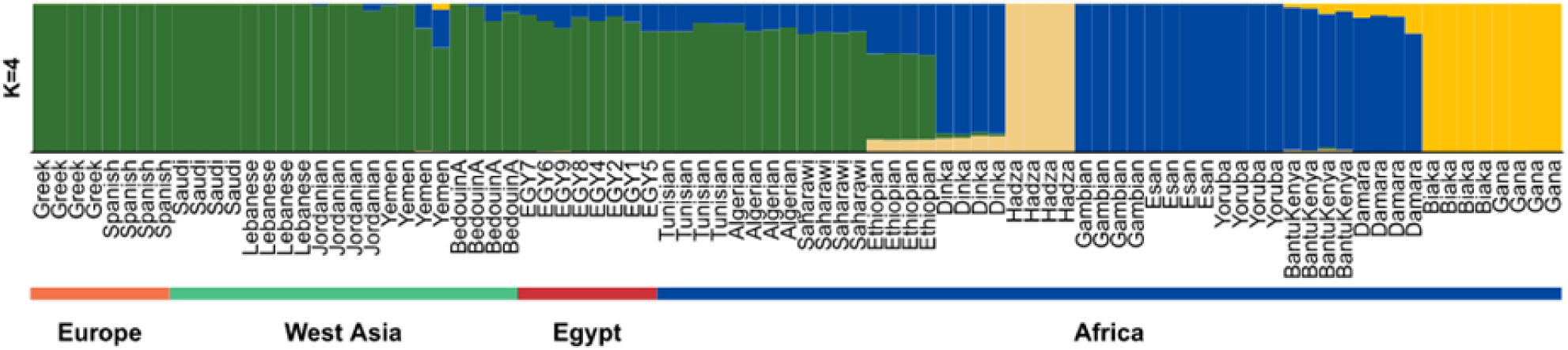
The ancestral composition of Egyptians in comparison to other populations when K = 4

## 4. Discussion

There is a limited amount of studies reporting complete genomes using NGS technology from different ethnic groups in North Africa [19,64,65,66], especially, together with clinical data. We detected the variants of the candidate genes, *GCAT, SLC6A13, TESK2, VCP, PCNX*, and *NUP88* using the CRAVAT v.4 server. Also, we found four already reported variants in *CRHR1* among the subjects who are suffering from severe obesity and metabolic syndrome. Rankinen *et al.* [67] demonstrated that *CRHR1* caused human obesity by single-gene mutations. [68,69]. Moreover, the participants suffering from obesity and obesity related diseases, (EGY2, EGY6, EGY7, and EGY9) had a SNP rs3733402, 428G>A (Ser143Asn) in *KLKB1*. This amino acid change has been associated with a Prekallikrein deficiency [70]. Also, we found Gly388Arg in *FGFR4* gene (rs351855, 1162G>A) in the participants EGY1, EGY6, and EGY8, which is associated with Waist-to-hip ratio adjusted for body mass index [36]. Moreover, four SNPs were found in pre-diabetic subject with HOMA-IR >2.5 (EGY6), and two diabetic subjects (EGY7, EGY9). One of these SNPs was in *MEFV*, 2080A>G, Met694Val, rs61752717 that is associated with Familial Mediterranean fever [71]. It is reported that *MEFV* gene variations affect mostly populations living in the Mediterranean region, especially North African Jews, Armenians, Turks, and Arabs and were also reported in a sequenced Bedouin genome[72].

The mtDNA analysis in the presented study revealed haplogroup for each subject as well as pathogenic mutations that can be transferred from mother to children. A mtDNA T16189C mutation observed in EGY7, EGY9, and EGY2, is already linked with insulin resistance and type 2 diabetes mellitus from the pooled case-manipulate research comparing the linkage of T16189C polymorphism with the threat of most cancers and T2DM progression [73]. Regarding other pathogenic mutations, each subject of the nine samples shared mutation A4282G (Table 2) related to chronic progressive external ophthalmoplegia (CPEO Plus) which is a slowly progressing disease that may begin at any age and progresses for fifteen years and often manifests in older age [74].

Overall, both mtDNA and ADMIXTURE studies suggested a diverse and admixed background of Egyptians sharing genetic signature and metabolic phenotypes with other Mediterranean nations. Although largely debated, into-Africa migration hypothesis [75] [76,77] seems to be supported by our largely diverse mtDNA haplotype collection demonstrated by just nine random samples. Egypt standing at the crossroad of different continents could have served as a gateway that allowed circulating people from Eurasia towards Africa since at least Holoscene [75]and lead to admixtures in east and sub-Saharan Africa that were reported before [76,77] and in this case lead to shared genetic predisposition to complex diseases such as obesity.

Our data can be a useful additional genomics resource for the greater genomic research of the nothern Africa and Middle East regions. The haplotype and admixture results point to the diversity of Egyptians, and to a non-African origin for most of them, pointing to the into Africa influx theory.

## 5. Conclusions

Here, we provide nine additional Egyptian genomic resources in terms of short-read based whole genome data and SNPs linked to metabolism-related phenotypes. These nine genomes are a valuable national resource as it contains matching clinical information for each of the nine participants. It can facilitate further larger-scale in depth research yielding more relevant variants and greater statistical power.

## Data

The datasets generated in this study are available in the NCBI Sequence Read Archive repository.

## Authors’ contributions

Idea and designed the experiments: MH. Performed the experiments: AE. Sequencing of the whole genomes: SS, TH, AK. Analyzed the data: EH, YJ, SJ, JB, AB. Wrote the paper: AE, EH, AB, JB. Study design, subject recruitment, and sample preparation: AE, FM. Data interpretation: AE, EH, SJ. Contribution to the final draft: AE, ME, SK, AB, SJ, YJ, JB. All authors read and approved the final manuscript.

## Acknowledgements

We thank the National Research Centre clinic for providing for the samples. We thank the pan Asian population genomics initiative (PAPGI) for support with sequencing. This work was supported by U-K BRAND Research Fund (1.190007.01) of UNIST; Research Project Funded by Ulsan City Research Fund (1.190033.01) of UNIST.

